# Symbiosis collapses during development of asexual offspring in the absence of heterotrophic feeding in a model cnidarian-algal symbiosis

**DOI:** 10.1101/2024.11.15.623823

**Authors:** Samuel A. Bedgood, Keyla Plichon, Virginia M. Weis

**Author notes:** Corresponding author: Samuel A. Bedgood.

## Abstract

A stable symbiosis between corals and dinoflagellate algae is crucial for coral reef health, and it is driven by nutrient exchange and environmental interactions. Our understanding of the homeostasis between host cnidarian and algal symbiont during the host adult stage is a longtime area of focus, but little is known about the balance of partners during development and regeneration. We investigated the role of symbiotic algae and heterotrophic feeding on development in the sea anemone model organism commonly called Aiptasia. We focused on asexually-produced offspring (G1), examining the effects of autotrophic and heterotrophic nutrition on developmental rates. We found that the presence of symbionts enhanced growth in fed conditions but impeded development and survival under starvation. The effect of symbiont presence on starved offspring was dose-dependent where those offspring with more symbionts at an earlier stage lost tentacles and mass faster than those with fewer symbionts. Our data demonstrate the importance of heterotrophic nutrition during early development and establishment of symbiosis. We propose that suppression of immunity during development may account for the observed patterns, although further research is required to validate this hypothesis. Our results provide insight into the metabolic costs and benefits of symbiosis under different nutritional conditions during development and regeneration of symbiotic cnidarians.

## 1. Introduction

Symbiosis between corals and dinoflagellate algae (Family Symbiodiniaceae) is foundational to a healthy coral reef ecosystem, and this relationship is based on nutrient exchange between partners^1–3^. The checks and balances to this relationship are complex and only partly understood. The homeostasis between the host (coral) and its symbiont (algae) regulates the biomass of each through three major processes: (1) nutrient dynamics, where the host controls nitrogen availability to the algae while symbionts supply photosynthetically derived sugars and lipids to the host^4–6;^ (2) host immune regulation of symbionts^8,9^; (3) symbiont population dynamics and cell cycle regulation in host tissue^7^; and (4) environmental factors such as light and temperature, which impact the host’s ability to maintain homeostasis and the symbiont’s survival and growth^10–12^. Changes in one or more of these factors can result in a move from symbiosis to dysbiosis whereby algae are lost from the host resulting in coral bleaching^13^. Environmental perturbations, due to climate change, are occurring on a global scale causing the degradation of coral reefs through breakdown in symbiosis^14^. It is therefore critical to understand the functioning of this symbiosis under different stressors and life stages of the host.

Cnidarians rely on a combination of autotrophic and heterotrophic nutrition to support their growth and metabolism. Algal symbionts provide photosynthetically derived carbon in the form of sugars and lipids, which fuel the host’s energy demands^15,16^. However, heterotrophic feeding on plankton and dissolved organic matter supplies essential nutrients, such as nitrogen and phosphorus^17,18^. This dual nutritional strategy reflects a trade-off, where autotrophy provides a consistent energy source while heterotrophy supplies essential nutrients, leading to variations in energy allocation depending on environmental conditions and resource availability^18,19^.

The influence of the host’s developmental stage on cnidarian-algae homeostasis remains understudied. Research in this area has primarily focused on how algal symbionts affect coral larvae energetics^20–23^, while asexually-produced offspring—an important component of most cnidarian life history strategies^24,25^—have largely been overlooked. Reef-building corals produce massive colonies via asexual polyp budding, colony fragmentation^26^, and asexual viviparity^27^.

Homeostasis between host and symbiont must be maintained throughout the process with the parent polyp(s) passing algae to the asexual offspring.

*Exaiptasia diaphana*, a sea anemone commonly known as Aiptasia, is a well-established model for the study of coral-algal symbiosis^28^. It reproduces both sexually and asexually. Sexual reproduction occurs through broadcast spawning of gametes, resulting in planula larvae that settle and develop into polyps^29–31^. These F1 offspring produced by F0 parents acquire their symbionts horizontally by ingesting algae during the planula or juvenile polyp stage^31^ (Fig. 1). Asexual reproduction occurs through pedal laceration, where adult anemones, referred to here as G0, leave behind small pieces of tissue from the pedal disk that develop into G1 offspring^32,33^.

**Figure 1.**
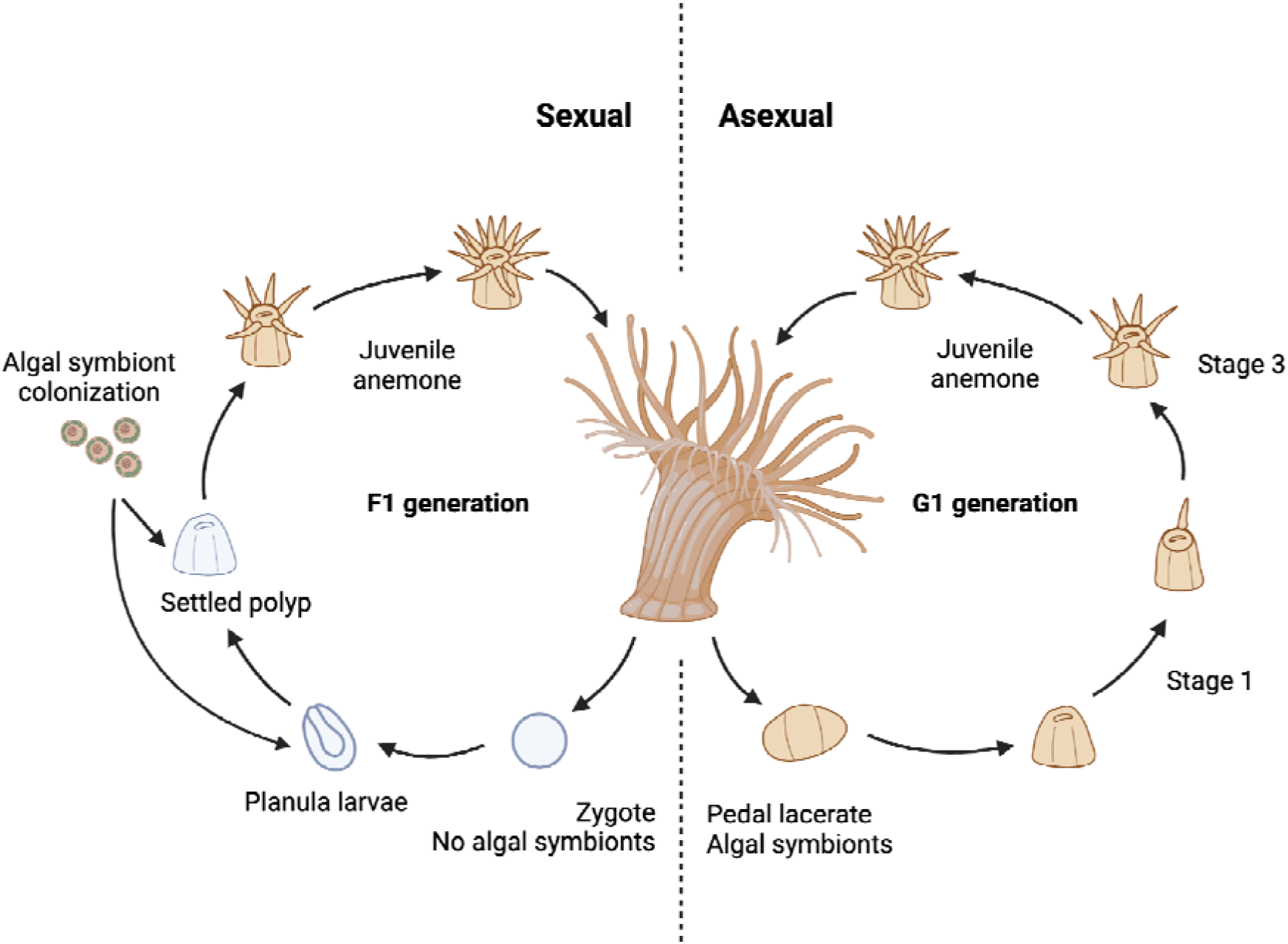
Reproductive modes of Aiptasia showing both the F1 (sexual reproduction) and G1 generation (asexual reproduction). Brown indicates the presence of algal symbionts and gray indicates no algal symbionts. Developmental stages are from Presnell et al. 2022. Created with BioRender.

Unlike F1 offspring, G1 offspring inherit their symbionts vertically, directly from the G0 adult^32^ (Fig. 1). Each pedal lacerate is on average 0.5 mm in diameter^33^ (S. Bedgood *pers. obs.*), but contains hundreds of algal cells from the parent (*pers. obs.*, S. Bedgood), which make up as much as 55-60% of the dry mass of a pedal lacerate^32^. Aposymbiotic pedal lacerates, like aposymbiotic adults, can acquire symbionts from the environment, once they have developed a mouth.

Presnell et al.^33^ showed that the presence of symbionts affects the developmental rate of G1 offspring when starved; aposymbiotic pedal lacerates develop more rapidly than symbiotic in the first few days of development. However, the growth rate in symbiotic offspring overtakes that of aposymbiotic offspring 20 days after separation from G0 anemones. The symbiosis becomes beneficial only after an initial dampening of growth rate. The role of heterotrophic feeding as an alternative or additive source of nutrition and the role of timing and onset of symbiosis have not been explored in these early developmental stages or in later stages of development beyond three weeks.

In this study, we extended the timeframe beyond the three-week window to explore the impact of nutrition, both from symbiotic photosynthate and heterotrophic feeding, on developmental rates. We also introduced the variable of horizontal symbiont transmission following laceration to determine effects from symbionts and nutrition during early-and late-stage development. We hypothesized that starved symbiotic G1 juveniles would exhibit a more rapid growth rate compared to their aposymbiotic counterparts in a timeframe longer than Presnell et al.^33^, suggesting that the symbiotic state confers a growth advantage under nutrient-limited conditions. Furthermore, we anticipated that starved, freshly inoculated G1 juveniles would demonstrate an intermediate growth rate: slower than symbiotic animals but faster than aposymbiotic animals. Lastly, we hypothesized that heterotrophic feeding would have an additive effect on growth rate, such that the growth benefit of feeding would be independent of symbiont presence, with both factors contributing separately to overall growth. Surprisingly, our results strikingly refute some of our hypotheses and bring up interesting new questions about the interacting effects of host age, nutrition, and early immunity on symbiosis health and regulation.

## 2. Methods

### ***a)*** Anemone and algal stocks

All sea anemone and algal cultures were kept in incubators (Percival Model: CU-36L5) at 25° C under 20–30 µmol quanta m^-2^ s^-1^ using 5,000K fluorescent lights (Zoo Med Flora Sun) on a 12/12h light/dark photoperiod. The algal cultures were maintained in 50 mL flasks using autoclaved F/2 media (Bigelow NCMA, East Boothbay, ME, USA) without silicates. New cultures were created once every two months by inoculating new F/2 media flasks with 5 mL of the previous culture. Sea anemones for this study came from the Aiptasia clonal line H2, originating from Hawaii^34^. To generate aposymbiotic (without algae) anemones, animals were artificially bleached with menthol^35^ once every three months. Aposymbiotic anemones used in this study were menthol-bleached two months before the experiment began to minimize effects from menthol on our results. Symbiotic parent anemones (G0s) were kept in clear 300 mL tubs containing 1 μm filtered seawater (FSW) while aposymbiotic animals were kept under the same conditions but in opaque containers to discourage re-inoculation with algae. G0 anemones were fed *ad libitum* three times weekly with freshly-hatched *Artemia* nauplii and 100% water changes with FSW were completed within 24 hours after feeding.

To verify the identity of algal species, algae from both G0 animals and algae in culture were sequenced using Internal Transcribed Spacer 2 (ITS2). DNA was extracted from whole anemones and algae pellets using the Qiagen DNeasy Plant Pro Kit. PCR amplification of the target region was performed with ITS2-specific primers (primer sequence). We analyzed the resulting PCR products through gel electrophoresis, followed by Sanger sequencing at the Center for Quantitative Life Sciences, Oregon State University. Sequences were cleaned and Symbiodiniaceae algae types identified using Geneious software and the SymPortal BLAST database^36^ to identify Symbiodiniaceae algae types. Algae from anemones and culture matched type B1, *Breviolum minutum*, the native symbiont to the H2 clonal line^37^. The algae within symbiotic parental sea anemones are assumed to be from the original clonal source while the culture was strain SSB01, an axenic culture sourced from Aiptasia^34^.

### ***b)*** Experimental design and husbandry

We prepared artificially-produced pedal lacerates (G1s) following Presnell et al.^33^ with some modifications. We placed symbiotic and aposymbiotic G0 adults in separate polypropylene petri dishes for two days so that they would fully attach. Once attached, we excised two pieces of tissue from the pedal disk of each anemone using a scalpel, for a total of 144 lacerates. The size of these artificially-produced lacerates was measured and the average size was not different from naturally-produced lacerates (data not shown). Each lacerate was placed in a well of a 12-well plate with 750 μL filtered seawater (FSW). The lacerates were left to develop naturally, not removing the mucous film that develops on the tissue ball as described in Presnell et al.^33^. We found no evidence for higher survival in lacerates that received this treatment (see Supp. Fig. 1).

We produced symbiotic state treatments: aposymbiotic, inoculated and symbiotic as illustrated in Figure 2. Symbiotic G1s were cut from symbiotic G0s, while inoculated and aposymbiotic pedal lacerates G1s were cut from aposymbiotic G0s. The inoculated treatment was colonized with algae (SSB01, *B. minutum*) in FSW at a final density of 3×10^5^ cells/mL within each well and left for 24 hours before a 100% water change. We inoculated using this method twice to ensure a successful inoculation, at four and then seven days after laceration. By day four, all anemones had developed a mouth, but only some had one tentacle (data not shown). By day seven, most anemones had six or more tentacles, following the pattern reported by Presnell et al.^33^. On day eight, we removed anemones from wells, the wells were cleaned with a cotton swab, and then the animals returned to the wells with fresh FSW. This removed any residual symbionts and mucus left in the wells from the first seven days. All inoculated anemones were confirmed to have algae by day 10 (data not shown).

**Figure 2.**
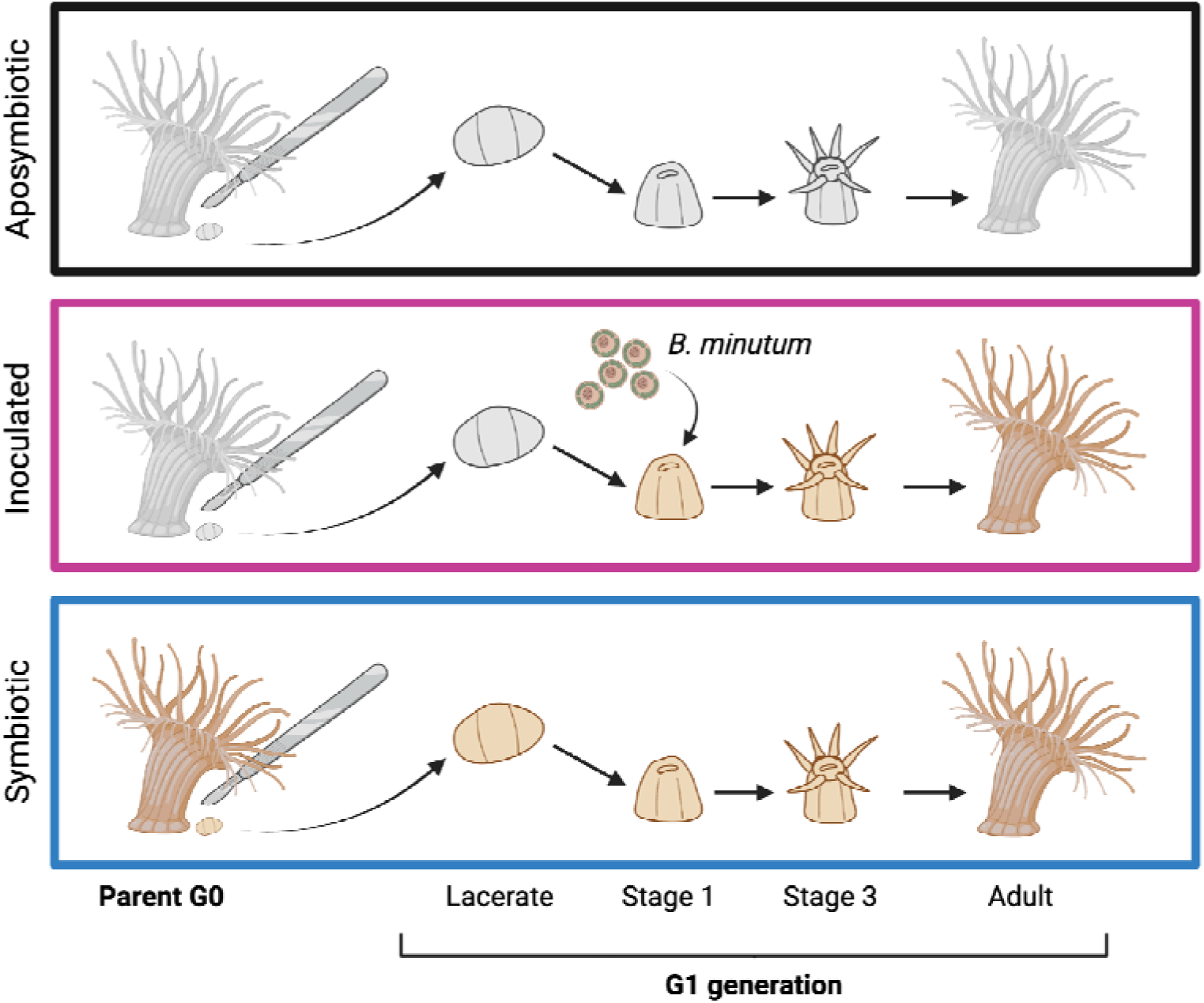
Illustration of the three algal symbiont experimental treatments. Brown indicates the presence of algal symbionts and gray indicates no algal symbionts. Developmental stages are from Presnell et al. 2022. Created with BioRender.

Symbiotic state treatments were crossed with two feeding treatments, starved and fed, for a total of 24 G1s in each treatment group at day zero. The starved treatments received no food and fed treatments were given 4-5 live rotifers (source culture from K&P Aquaculture) three times weekly beginning seven days after laceration. FSW changes were completed three times weekly, 24 hours after feeding. We quantified tentacle number five to seven times weekly beginning on day four. Tentacles begin as thick nubs and extend. We counted tentacles only if they were longer than they were wide. The rotifers fed to anemones were kept at 26°C in 1L bottles that received water changes once weekly. We sieved the rotifer culture with a 50 μm nylon mesh, placed the sieved rotifers back in the bottle, and replaced the water with 900 mL FSW and 100 mL *Tetraselmis* sp. microalgae once weekly. Live *Tetraselmis* sp. cultures were maintained in 1L bottles with F/2 media in FSW refreshed weekly under 200 µmol quanta m^-2^ s^-1^ LED lights (Barrina T5 Grow Lights, pinkish white).

### ***c)*** Symbiont density and size

We imaged G1 anemones on a Zeiss Axio Observer.A1 inverted epifluorescence microscope under brightfield illumination for pedal disk area and at 700 nm excitation for symbiont imaging. All images were analyzed in ImageJ^38^. Pedal disk area was used as a non-lethal measure of anemone size^39^. We measured symbiont density from images of tentacles by selecting a region in the middle of each tentacle 2-8 mm in length and divided the proportion of algae cell pixels (fluorescing) by the total number of pixels in the selected region, similar to Maruyama et al.^40^. We took the symbiont density of two tentacles per anemone. This allowed for multiple non-destructive measurements from the same individuals.

### ***d)*** Cell division

EdU (5-ethynyl 2’-deoxyuridine), a thymidine analog, gives a snapshot of host anemone cell division rates *in vivo* (proportion of cells in S-phase) within an incubation period. We sacrificed four anemones from each treatment for EdU labeling at three time points, after the second, fourth, and tenth week. By the tenth week, there were no starved symbiotic or starved inoculated G1s to sample. G1 anemones were incubated in 10 μM EdU in their wells for 24 hours before fixing in 4% formaldehyde, 0.2% glutaraldehyde, 0.1% Tween in PBS for 1.5 minutes then 4% formaldehyde and 0.1% Tween in PBS overnight. Each polyp was treated as described in Tivey et al.^7^ to label EdU+ cells (Click-iT EdU Alexa Fluor 555 imaging kit; Life Technologies, Eugene, OR, USA) and host nuclei with Hoechst 33342. Briefly, tissue was permeabilized in 0.2% PBST (0.2% Triton in PBS), washed in 0.02% PBST, blocked in 5% BSA in PBS, labeled with EdU reaction, rinsed with 0.02% PBST, labeled with Hoechst 33342, washed in 0.02% PBST, and mounted on a slide suspended in a final concentration of 87% glycerol. Slides were imaged on a Zeiss LSM 780 NLO confocal microscope with three excitation wavelengths: 555 nm (EdU-AF555), 405 nm (Hoechst), and 633 nm for algal symbiont autofluorescence. Images were taken at multiple depths and combined into z-stacks for better resolution across three dimensional structures. These z-stack images were analyzed in FIJI^38^ by selecting a region in the middle of a tentacle 2-8 mm in length and measuring the number of pixels of EdU fluorescence standardized by the number of pixels of Hoechst fluorescence to calculate a host cell division rate.

### ***e)*** Statistical analyses

All analyses were completed in R v. 4.1.2^41^. We used the package lme4 to create models^42^, car for ANOVA^43^, and emmeans for posthoc analyses^44^. All models followed the same design, which included a three-way interaction among symbiotic state treatment, feeding treatment, and time, lm(data ∼ symbiont * feed * day). We analyzed tentacle count data with a generalized linear mixed model with a log link Poisson distribution. We analyzed symbiont density, host size, and host cell division rate with linear models and checked differences between observed and predicted values (residuals) for normality before analysis.

## 3. Results

### ***a)*** Survival of starved anemones was reduced by the presence of symbionts

All anemones survived the initial development period before day 30, except for three in the symbiotic treatments (Fig. 3). By day 30, the starved symbiotic anemones group had lost four individuals while the fed symbiotic anemones group had lost one out of 24. After day 30, the starved symbiotic and inoculated anemones rapidly died until there was only one G1 left in the inoculated treatment. The rate of loss between starved symbiotic and inoculated anemones was similar, but the inoculated group lagged approximately one week behind the symbiotic group.

**Figure 3.**
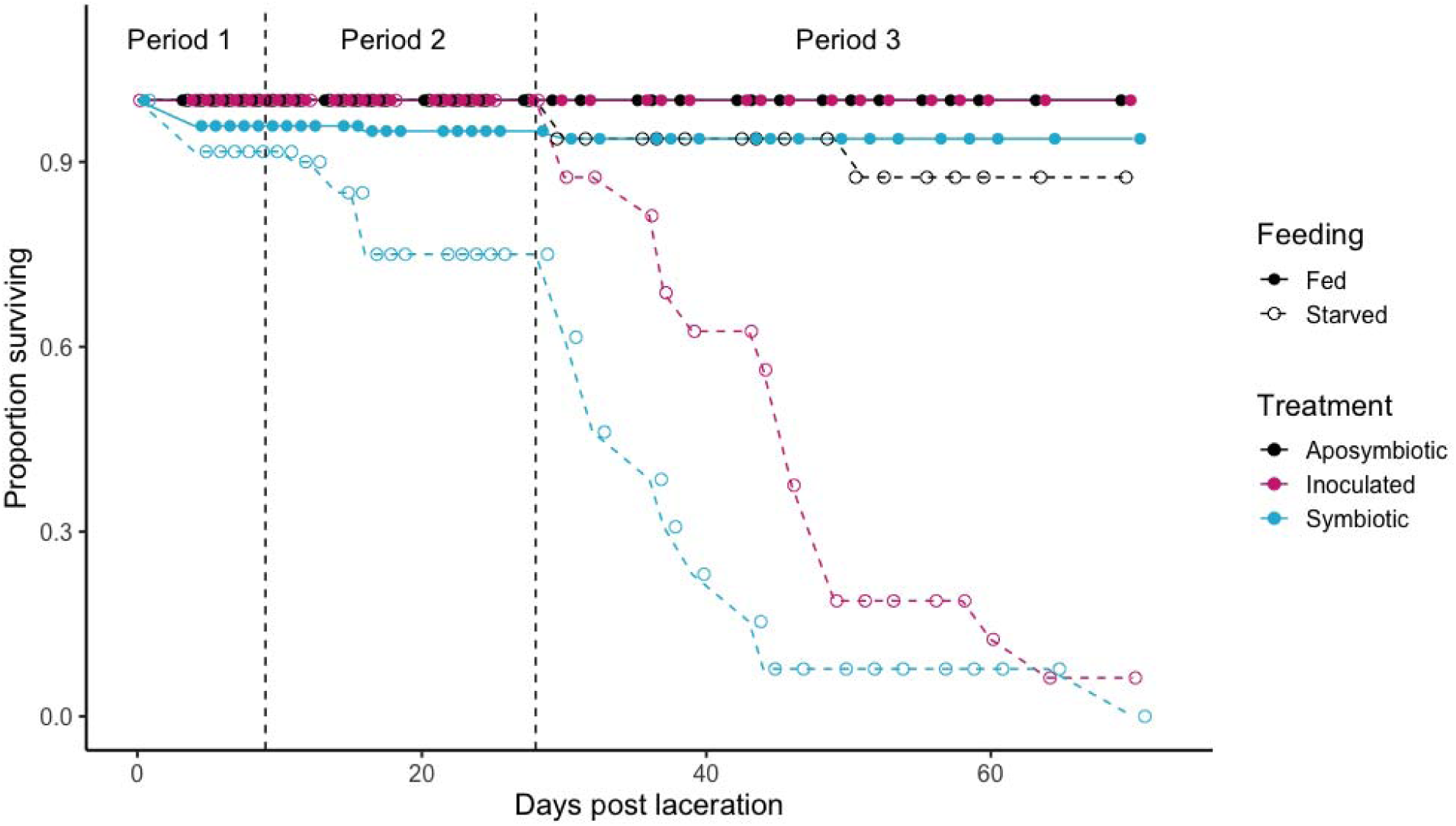
Proportion of pedal lacerates surviving over time (days post-laceration). Each treatment began with 24 individuals. Vertical dashed lines separate Periods in the experiment. Starved symbiotic offspring exhibited mortality starting in Period 2, while starved inoculated anemones began to die at the beginning of Period 3. By day 70, only one starved anemone with symbionts remained alive, with all others having died. Survival was highest in fed aposymbiotic and inoculated anemones.

Only two anemones died in the starved aposymbiotic treatment group by the end of the experiment, and no fed anemones died after day one.

### ***b)*** Development was affected by an interaction between feeding and the presence of symbionts

We identified three distinct developmental periods based on observed changes in the data: Period 1 (days 0-9), during which all lacerates developed eight tentacles; Period 2 (days 10-28), characterized by a divergence between fed and starved treatments; and Period 3 (days 29-70; Fig. 4), when starved symbiotic anemones began to die. During Period 1, symbiotic state had an effect on host development (symbiont: 2 = 77.38, p < 0.001; symbiont*day: 2 = 78.08, p < 0.001; Fig. 4) similar to what was found during the first few days of development by Presnell et al.^33^. Symbiotic G1s developed more slowly than all other anemones in other treatments, including G1s that were inoculated with algae on days four and seven. Symbiotic G1s had significantly fewer tentacles than all other groups on days 5-7 (Supp. Table 1). Feeding had a small effect on development during Period 1 where fed animals grew tentacles at a faster rate (□2 = 4.93, p = 0.026). However, symbiotic G1s caught up to animals in the rest of the treatments by day nine, the beginning of Period 2. There were no significant differences between treatment groups on day nine.

**Figure 4.**
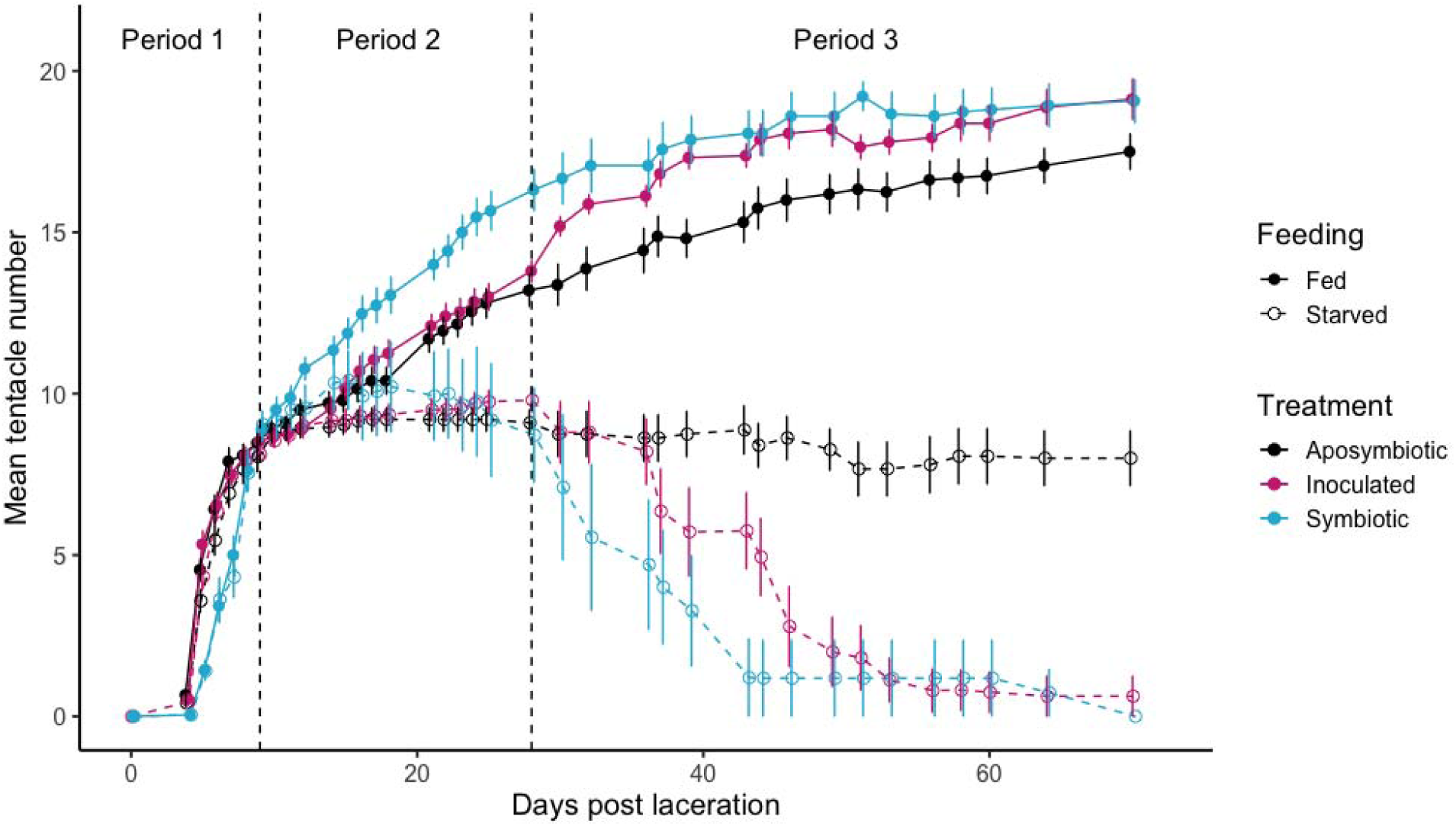
Pedal lacerates tentacle count over time (days post-laceration). Points are jittered to differentiate error bars. Points are means ± standard error of the mean. Vertical dashed lines separate Periods in the experiment. Sample sizes for each treatment at each time point were reduced throughout the experiment because of lethal sampling for EdU labeling where days 0-15 had N = 24, days 16-29 had N = 20, and days 30-70 had N = 16. During Period 1, symbiotic anemones exhibited a slower developmental rate compared to other groups. In Period 2, the influence of food availability became apparent, as fed anemones showed faster development. By Period 3, there was a clear interaction between symbiont presence and feeding; the presence of symbionts was only beneficial to development when food was available.

During Period 2, there was a strong interaction between feeding and symbiotic state (□2 = 10.44, p = 0.005) where fed anemones developed faster than starved ones and the presence of symbionts increased growth rates only when combined with feeding. (Fig. 4). Fed symbiotic anemones, overtook animals in the rest of the treatments with an average of 16.3±0.6 (mean±SEM) tentacles by day 28. All starved treatments performed similarly, with G1s in the starved symbiotic treatment having an average of 8.7±1.5 tentacles by the end of the period.

Tentacle numbers of fed symbiotic and fed inoculated animals were not significantly different (z =-2.02, p = 0.329) and fell between these two groups at an average of 13.8±0.3 tentacles in the fed inoculated treatment by day 28.

During Period 3, there was a strong interaction between feeding and symbiotic state treatments ( 2 = 948.0, p < 0.001) where the presence of symbionts, both in the symbiotic and inoculated treatments, had a positive effect on development if food was present but, crucially, a deleterious effect when the host anemone was starved (Fig. 4). G1s in the fed inoculated treatment caught up to those in the fed symbiotic treatment by the end of Period 3 (z = 0.03, p = 1.0). G1s in the fed aposymbiotic treatment had fewer tentacles than the fed symbiotic and inoculated G1s throughout this period (Fig. 4), but no pairwise comparisons were significant (Supp. Table 1). The starved aposymbiotic G1s had fewer tentacles than the fed aposymbiotic G1s (day 70: z = 0.12, p < 0.001), but more tentacles than starved symbiotic and inoculated G1s (day 64 starved symbiotic: z = 6.58, p < 0.001; day 64 starved inoculated: z = 7.76, p < 0.001).

The starved anemones that had algal symbionts notably lost tentacles during Period 3 (Fig. 4). G1s that were starved and had symbionts began losing tentacles on day 28 (Fig. 4). The starved symbiotic G1s lost tentacles earlier than the starved inoculated G1s, but the starved inoculated anemones caught up to the loss three weeks later (Fig. 4). Most starved anemones that had symbionts lost all of their tentacles, eventually forming balls of tissue that were dead by day 70.

### ***c)*** Symbiont density was higher in starved anemones

There was an effect of feeding (F = 56.34, p < 0.001), symbiotic state (F = 122.55, p < 0.001), and an interaction between feeding and symbiotic state (F = 18.38, p < 0.001) on symbiont density in host tissues. The inoculated G1s began with no symbionts at day zero while the natural-state symbiotic G1s started with hundreds of symbionts from G0s (*pers. obs.*, S. Bedgood). Symbiont density was first measured at day 10 when well-developed tentacles were present. By this time, the starved symbiotic G1s had a higher symbiont density than the fed symbiotic G1s (t =-6.31, p < 0.001), and this pattern persisted throughout the experiment (Fig. 5). The symbiont density within the inoculated anemones was lower than the symbiotic anemones at day 10 (Fig. 5), and there was no difference between fed inoculated and starved inoculated on day 10 (t =-0.05, p = 0.999) or day 25 (t =-1.70, p = 0.329). However, by day 25 the inoculated G1s had a similar density to the symbiotic G1s (Fig. 5). On day 66 there was only one anemone left in the fed symbiotic group, but this anemone had 100% symbiont fluorescence per tentacle area (Fig. 5). The starved inoculated G1s had a higher symbiont density than the G1s in either of the fed treatments (fed symbiotic t = 6.27, p < 0.001; fed inoculated t =-6.55, p < 0.001), and there was no difference between the fed symbiotic and fed inoculated anemones (t =-0.37, p = 0.982), which had the lowest density (Fig. 5). No algae was detected in the aposymbiotic anemones throughout the experiment.

**Figure 5.**
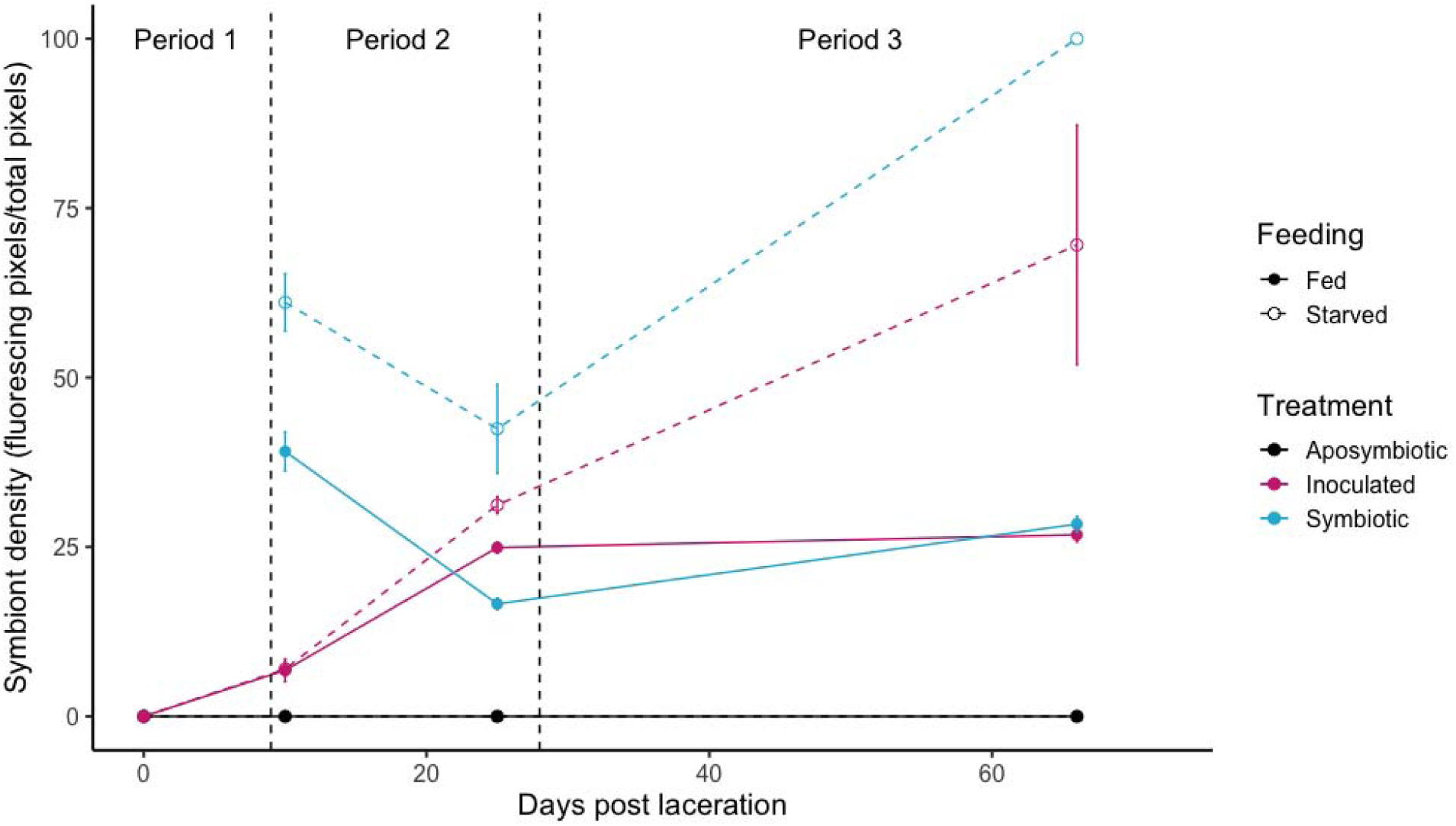
Symbiont density of pedal lacerates over time (days post-laceration). The symbiont density of symbiotic lacerates at time point zero was not measured. Points are means ± standard error of the mean. Vertical dashed lines separate Periods in the experiment. Sample sizes for each treatment at each time point were reduced throughout the experiment because of lethal sampling for EdU labeling where days 0-15 had N = 24, days 16-29 had N = 20, and days 30-70 had N = 16. Symbiont density began higher in symbiotic anemones compared to inoculated anemones. Throughout the experiment, fed anemones with symbionts maintained a stable symbiont density, while starved anemones showed a notable increase in symbiont density, reaching more than double the density observed in the fed treatment.

### ***d)*** Growth was increased by symbionts only in the fed anemones

Size of anemones was affected by symbiotic state (F = 68.19, p < 0.001), feeding (F = 123.91, p < 0.001), and a three-way interaction between symbiotic state, feeding, and time (F = 7.15, p < 0.001). We visually size matched lacerates while cutting them for our treatments, but by day two the symbiotic lacerates were significantly larger than all other treatments (Fig. 6; Supp. Table 3). On days 10 and 24, the inoculated G1s had similar sizes to their respective aposymbiotic treatments (Fig. 6; Supp. Table 3). On day 66 there was a clear interaction between symbiotic state and feeding. There were no differences between any of the starved anemone groups (Supp. Table 3) which lost 86.65±0.28% of their initial size, but there were large differences among the three fed anemone groups. The fed symbiotic anemones were larger than the fed inoculated anemones (t =-3.43, p = 0.009), and the fed inoculated anemones were larger than the fed aposymbiotic anemones (t =-5.70, p < 0.001).

**Figure 6.**
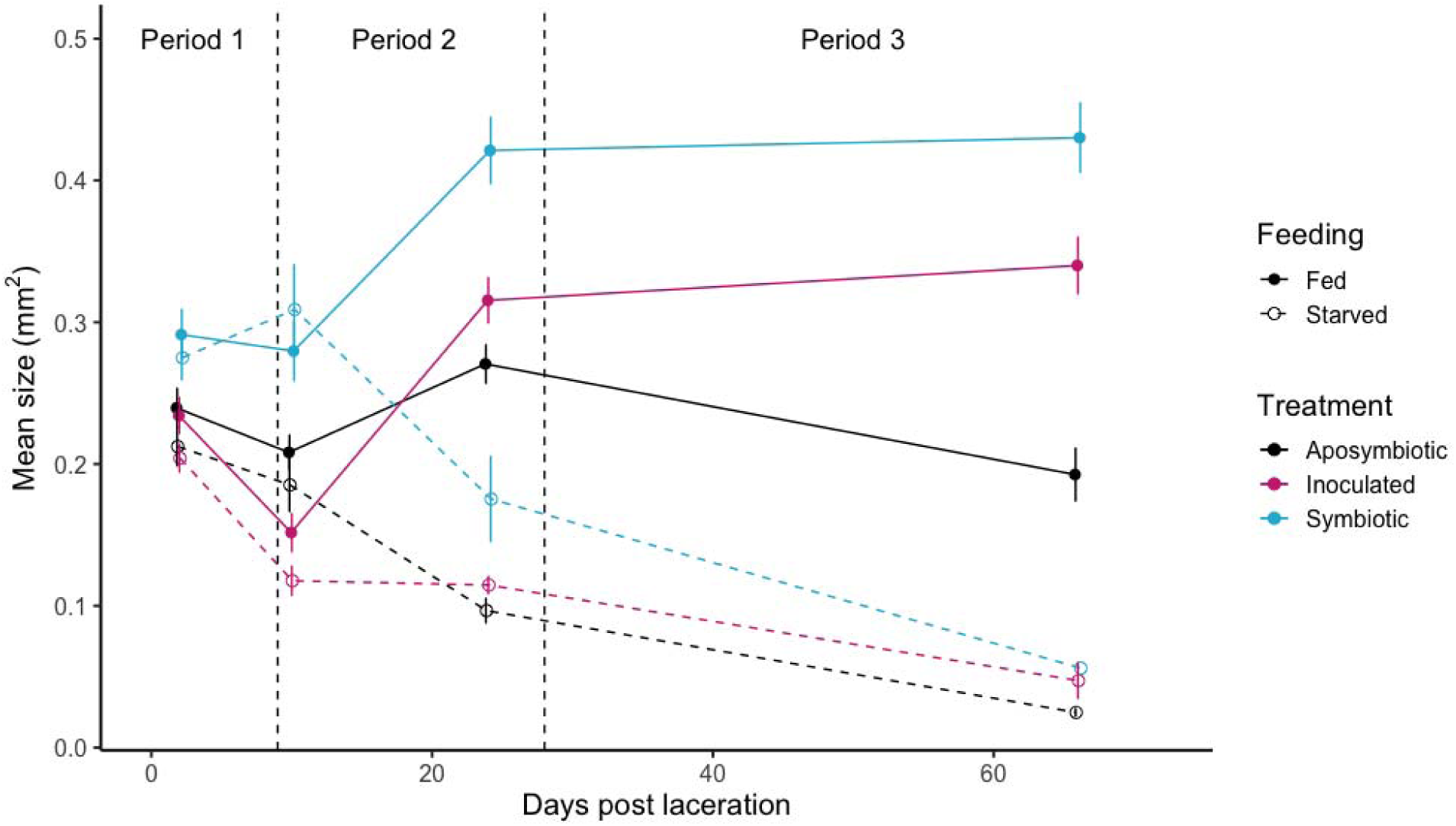
Size of pedal lacerates over time (days post laceration). Points are means ± standard error of the mean. Vertical dashed lines separate Periods in the experiment. Sample sizes for each treatment at each time point were reduced throughout the experiment because of lethal sampling for EdU labeling where days 0-15 had N = 24, days 16-29 had N = 20, and days 30-70 had N = 16. Symbiotic lacerates were larger during Period 1, but those groups that were fed grew larger during Periods 2 and 3. The presence of symbionts increased size only in the fed groups.

### ***e)*** Host cell division was higher in fed anemones

Host cell division rate, as measured by EdU fluorescence was affected by feeding (F = 10.70, p = 0.002) but unaffected by symbiotic state (F = 18.46, p < 0.001). There were some pairwise differences between symbiotic states on day 15 where fed symbiotic animals had higher cell division than fed inoculated (t =-3.20, p = 0.030) and fed aposymbiotic anemones (t = - 3.22, p = 0.028), but the effect of symbiotic state in the fed anemones was gone by the next time point (aposymbiotic vs. symbiotic: t =-0.722, p = 0.978). Fed anemones had higher cell division than starved aposymbiotic anemones on days 29 and 70 (Supp. Fig. 2, Supp. Table 4).

## 4. Discussion

Our study demonstrated an interaction between symbiotic state and feeding on survival, development, and growth of G1 (asexually-produced) offspring in a model cnidarian-algal symbiosis. There was a strong interaction between symbiotic state and feeding where symbionts were beneficial to the developing host anemone only when the host was fed. Fed anemones with algae developed faster and grew larger than those without algae, and this effect was symbiont dose-dependent. G1 anemones that started with symbionts grew faster than those that were inoculated later and only started with a few cells that had to proliferate throughout the host. This supported our initial hypothesis that the presence of symbionts would increase the growth rate of G1 offspring.

The presence of symbionts in the starved treatments, however, did not increase the growth rate of anemones, and, instead, slowed and eventually reversed tentacle growth after the first three weeks, possibly after the offspring exhausted the resources from G0 adults. Starved anemones shrank regardless of symbiotic state, but those with algae were more likely to lose all tentacles and die. The effect was more pronounced in anemones with higher initial symbiont densities. Symbiont density increased seemingly unchecked; all starved anemones with symbionts met the same fate: losing tentacles and eventually becoming balls of host tissue full of algal cells. This contradicted our hypothesis that the presence of symbionts would boost development and growth. Instead, high symbiont densities in starved G1 anemones likely exacerbated the adverse effects of starvation leading to their demise, as visualized by images of EdU labeled dividing cells in fed compared to starved animals (Fig. 7).

**Figure 7.**
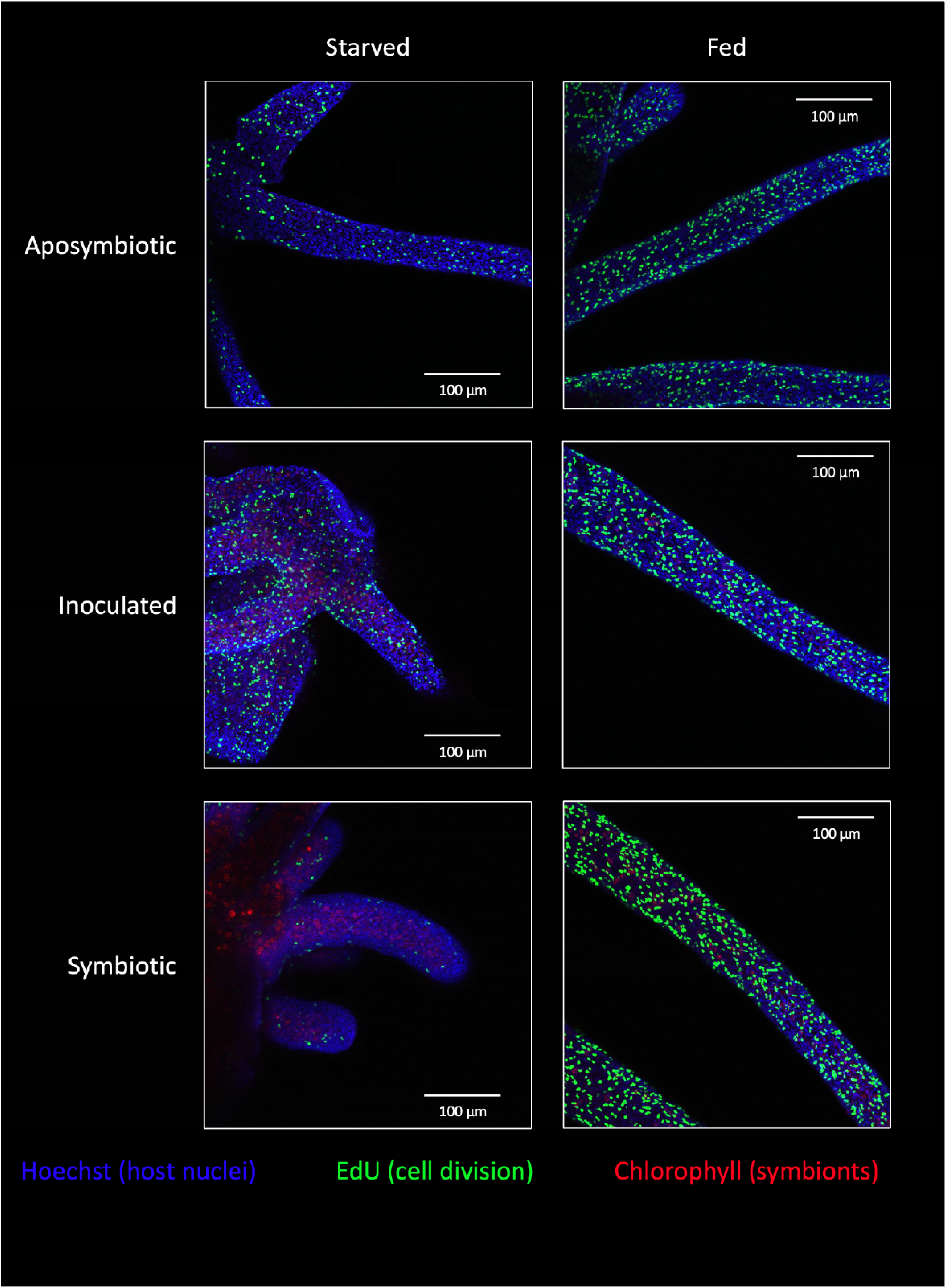
Representative z-stack images of anemones from day 29 (beginning of Period 3) of the experiment incubated in EdU for 24 hours before fixation. Fluorescent channels mark host nuclei in blue (Hoechst), host cell division in green (EdU), and algal symbionts in red (chlorophyll autofluorescence).

Our findings expand on previous studies on sea anemones and corals. The symbiosis-by-nutrition interaction has recently been a subject of studies performed in both adult corals^45^ and Aiptasia^46^. When corals were deprived of nutrients, for many months, they were shown to ‘farm’ their algae: digest a portion of the algal population to help them survive low nutrient conditions^45^. When anemones were starved for one year, both host biomass and symbiont densities in host tissues decreased dramatically. However crucially, the symbionts continued to pass carbon to the host: symbiosis homeostasis was maintained, the animals remained symbiotic and the holobiont remained healthy^46^. These results are in marked contrast to those found in our G1 juveniles where starvation caused profound symbiosis dysregulation and led to death.

A study on Aiptasia by Peng et al.^46^ found that aposymbiotic anemones inoculated with algae and then starved for months demonstrated a fatal metabolic cost from algae, leading to high mortality rates in the inoculated anemones as compared to the aposymbiotic controls. Our results focusing on G1 offspring from pedal laceration, suggest that developmental stage plays a crucial role in the regulation of homeostasis of the symbiosis in the absence of heterotrophic feeding.

The key differences between our study and Peng et. al.^46^ is that we included symbiotic G1s from the start while Peng’s study included only aposymbiotic and inoculated G1s, and we followed G1 anemones from initial laceration to adults while Peng’s study started with juveniles (6 tentacles, Stage 3 juveniles^33^). Our results also align with a modeling study that used data from the Peng et al.^46^ study to develop a food-dependent model in Aiptasia^47^. The model developed in the study “demonstrates the role of the symbiont as an amplifier of the host’s state: a growing anemone grows better with symbionts, while a malnourished anemone loses biomass faster with a symbiont than without.” Our results recapitulate and expand on these studies, demonstrating a stronger metabolic cost of algal symbionts when algae are transmitted vertically at higher densities and at earlier developmental stages. This indicates that earlier developmental stages may be less capable of regulating symbiont populations under nutrient-limited conditions than adults, leading to symbiosis dysregulation and host mortality.

Taken together, these studies suggest that host-symbiont biomass regulation and homeostatic regulation of symbiosis differs between adult and developing animals. What processes could explain this pattern? We suggest that the differences may be influenced by two factors: 1) nitrogen-limitation within the holobiont and 2) host immunity.

The runaway symbiont density in starved G1 anemones may be a result of nitrogen limitation within the holobiont. Photosynthetically derived carbon (sugars and lipids) is provided by the algal symbionts, but nitrogen comes from heterotrophic feeding^19,48^. Nitrogen, in the form of ammonium, can be passed to algal symbionts or recycled by the host using organic carbon from algal symbionts to produce amino acids^5^. This framework suggests a competition between host and symbiont for heterotrophically-derived nitrogen^6^. A severe carbon-nitrogen imbalance may occur in those developing offspring where the host digests its own tissue to meet nitrogen demands^1^. In fact, a phenotype of Aiptasia that lacks tentacles “Wurst” has been described by Rädecker et al.^49^ and is induced by starvation of adult anemones. These nutrient deprived adults produce pedal lacerates that never develop tentacles.

Nitrogen limitation does not account for the differences observed between starved symbiotic adult anemones in Peng et al.^46^, which showed no shrinking of tissue, and our early-development offspring, which lost all tentacles. Host immunity and symbiosis reestablishment during development could explain the pattern. Asexual reproduction and regeneration in early-diverging metazoans are evolutionarily tightly linked. Indeed, pedal laceration can be considered a form of whole-body regeneration (WBR), an area of evolutionary developmental biology that has been of interest to biomedicine for many years^50^. In Aiptasia, regeneration, including pedal laceration, can trigger immune responses that are linked with developmental pathways^51^. Studies on WBR in a variety of animals including cnidarians suggest an inverse relationship between regenerative/developmental processes and immune activation^52,53^. Transcriptomic studies on Aiptasia regeneration have shown that innate immunity pathways are initially upregulated following wounding but are downregulated as developmental programs take over^51^. This could explain our results in G1 juveniles. After laceration, immunity is repressed to facilitate WBR. However, when heterotrophic feeding is withheld and with immunity downregulated, the symbiosis becomes dysregulated, leading to unchecked parasitic algal proliferation within the host tissue and a fatal cost to the developing host. In contrast, when food is present, dysregulation is avoided and normal development proceeds.

Host immune response can also be affected by symbiont density in cnidarians. A study on *Orbicella faveolata* has demonstrated that high symbiont densities, produced with high ammonium concentrations, can significantly suppress immune gene expression^54^. However, in *A. poculata*, a positive correlation between symbiont density and immune response was observed, and the immune response varied with symbiont density^55^. These findings suggest that symbiont density is a critical factor that can influence immunity, and dysregulation of algal symbionts can lead to adverse outcomes, such as those observed in our starved and symbiotic G1 anemones.

To further our understanding of the interplay between nutrition, development, and symbionts, future studies could focus on algal cell cycle regulation during development using methodologies similar to those employed by Tivey et al.^7^ and transcriptomic comparisons across different feeding conditions, developmental stages, and symbiotic states. Examining sexually-produced juvenile polyps, F1 offspring, could provide insights into whether the outcomes observed in G1 offspring are consistent across different reproductive modes. Maintaining homeostasis between partners in coral-algal symbiosis during development, regeneration, and growth is essential for healthy coral reefs. A focus on nutrition during early development and/or regeneration could be critical for successful conservation efforts.

## Data Availability Statement

The datasets generated and analyzed during this study are available in the Dryad repository at http://datadryad.org/stash/share/ERsl_MoJ_uTCmOJQJai99d2rPAwFKJeKBwv1g0n-iaU.

## Acknowledgements

We thank Ruby Miller and Felix Jardini for their assistance in carrying out the experiment, caring for the animals, and counting thousands of tentacles. We also thank Sofia Blaire for contributing to the project and conducting a parallel experiment that informed the design of this study. We appreciate the guidance of Dr. Nathan Kirk and Dr. Keith Garrison in sequencing.

Additionally, we thank the members of the Weis Lab for their valuable input on the experimental design and analysis. Finally, we acknowledge the Center for Quantitative Life Sciences at Oregon State University for their confocal microscopy and sequencing services, which were essential to this work.

This research was supported by the National Science Foundation (NSF) through a grant awarded to Virginia M. Weis (NSF Grant No. IOS2124119). The funding agency had no involvement in the study design, data collection, analysis, or interpretation of the results, nor in the decision to submit the manuscript for publication.

## Author Contributions

S.A.B. conceived and designed the study, conducted the experiments, performed statistical analyses, and wrote the manuscript. V.M.W. contributed to the study design and manuscript writing. K.P. assisted with experimental work and provided critical feedback on the manuscript. All authors reviewed and approved the final version of the manuscript.

## Additional Information

The authors declare no competing interests.

