## Supplementary material for "Symbiosis collapses during development of asexual offspring in the absence of heterotrophic feeding in a model cnidarian-algal symbiosis": Supp. Figures

**Supplemental Figures**

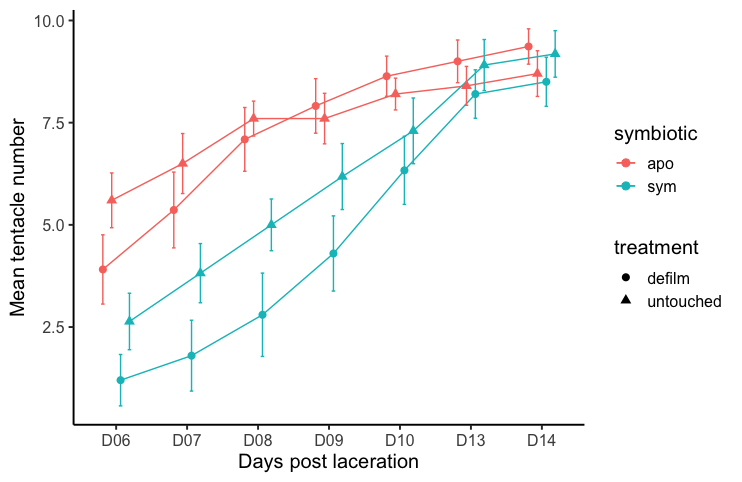

**Supplemental Figure 1.** Pedal lacerates tentacle count over time (days post-laceration). Points are jittered to differentiate error bars. Points are means ± standard error of the mean. Sample size was n = 10 for each treatment at each time point.

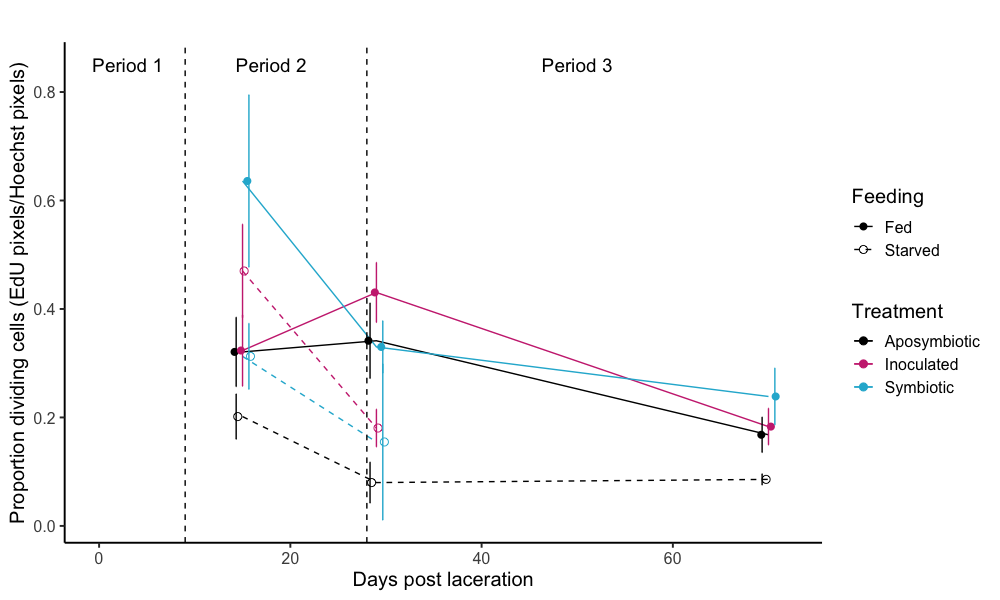

**Supplemental Figure 2.** Cell division (EdU fluorescence/Hoechst fluorescence) over time (days post laceration). All starved symbiotic and inoculated offspring died by the final time point (day 70). Points are means ± standard error of the mean.

**Supplemental Table 1.** Attached document labeled “Supp. Table 1.xlsx.” Posthoc pairwise analysis results from the tentacle count generalized linear model. Data are illustrated in Figure 4. Treatment groups were compared within time points (days).

**Supplemental Table 2.** Attached document labeled “Supp. Table 2.xlsx.” Posthoc pairwise analysis results from the symbiont density general linear model. Data are illustrated in Figure 5. Treatment groups were compared within time points (days).

**Supplemental Table 3.** Attached document labeled “Supp. Table 3.xlsx.” Posthoc pairwise analysis results from the anemone size general linear model. Data are illustrated in Figure 6. Treatment groups were compared within time points (days).

**Supplemental Table 4.** Attached document labeled “Supp. Table 4.xlsx.” Posthoc pairwise analysis results from the cell division (EdU) general linear model. Data are illustrated in Supplemental Figure 1. Treatment groups were compared within time points (days).
